# TE-aware state analysis and evolution in dynamic network connectivity

**DOI:** 10.64898/2026.09.20.753026

**Authors:** Micah Holness, Anastasia Bohsali, Brad Baker, Sir-Lord Wiafe, Zijing Dong, Fuyixue Wang, Lisa Krishnamurthy, Vince Calhoun

## Abstract

Multi-echo functional magnetic resonance imaging (fMRI) acquires signals with distinct contrast profiles across echo times (TEs). Empirical evidence suggests that different TEs capture distinct signal contributions with varying sensitivity to BOLD and non-BOLD processes from long to short TEs (Stroman, 2002; Krishnamurthy, 2023; Dong, 2024). Previous literature has not yet examined how brain states extracted from temporal dynamic functional connectivity (dFNC) evolve across TEs. Here we present a methodological and empirical study of TE-dependent evolution of dynamic connectivity states, with a phenomenological two-component model for interpretation. Multiple echoes were acquired with an echo planar time-resolved imaging (EPTI) sequence. TE-specific components were extracted via atlas-based group ICA (Neuromark 1.0), and brain connectivity states were generated via windowed dFNC and spatial ICA clustering of windows. Within-subject and group-level states evolved across TEs in network organization and variance. States exhibited TE-dependent variance profiles, with distinct peak echo times; and group-level points of state transition were independently mapped to TE-dependent transitions in variance. We modified a bi-exponential model to simulate and model changes in the dFNC time-course across TEs. The bi-exponential model revealed high consistency with subject states (R2 fit=0.86, res. error=0.55, 86% model convergence rate), suggesting a two-component signal system contributing to dynamic TE-dependent connectivity changes in state. A significant correlation between relative model terms was estimated on a cell-wise basis across states (*r*(24) = -0.43, *p*=0.03, 95% CI [-0.70, -0.04]). This suggests that specific bi-exponential signal parameters (i.e., short FNC_1,TE=0_, long FNC_2,TE=0_, and long FNC_2,decay_) were associated with the observed TE-dependent variations in connectivity states across TEs. In sum, our analyses provide downstream implications for TE-aware state analysis dependent on connectivity magnitude (FNC_TE=0_) and rate of connectivity decay (FNC_decay_), in a two-component signal model, with future implications for TE-dependent state biomarkers.

**Highlights:**

- Echo times are concatenated across this dimension, generating novel TE-aware dynamic changes in state connectivity.
- Bi-exponential fitting to dFNC signal evolution across TEs allows sensitivity to evaluate parameters involved in TE-dependent connectivity.
- TE-aware state analyses currently enhances multi-echo fMRI utility in a basic science context, with implications for TE-aware methodology to have valuable use in future clinical contexts.

**Graphical Abstract:** 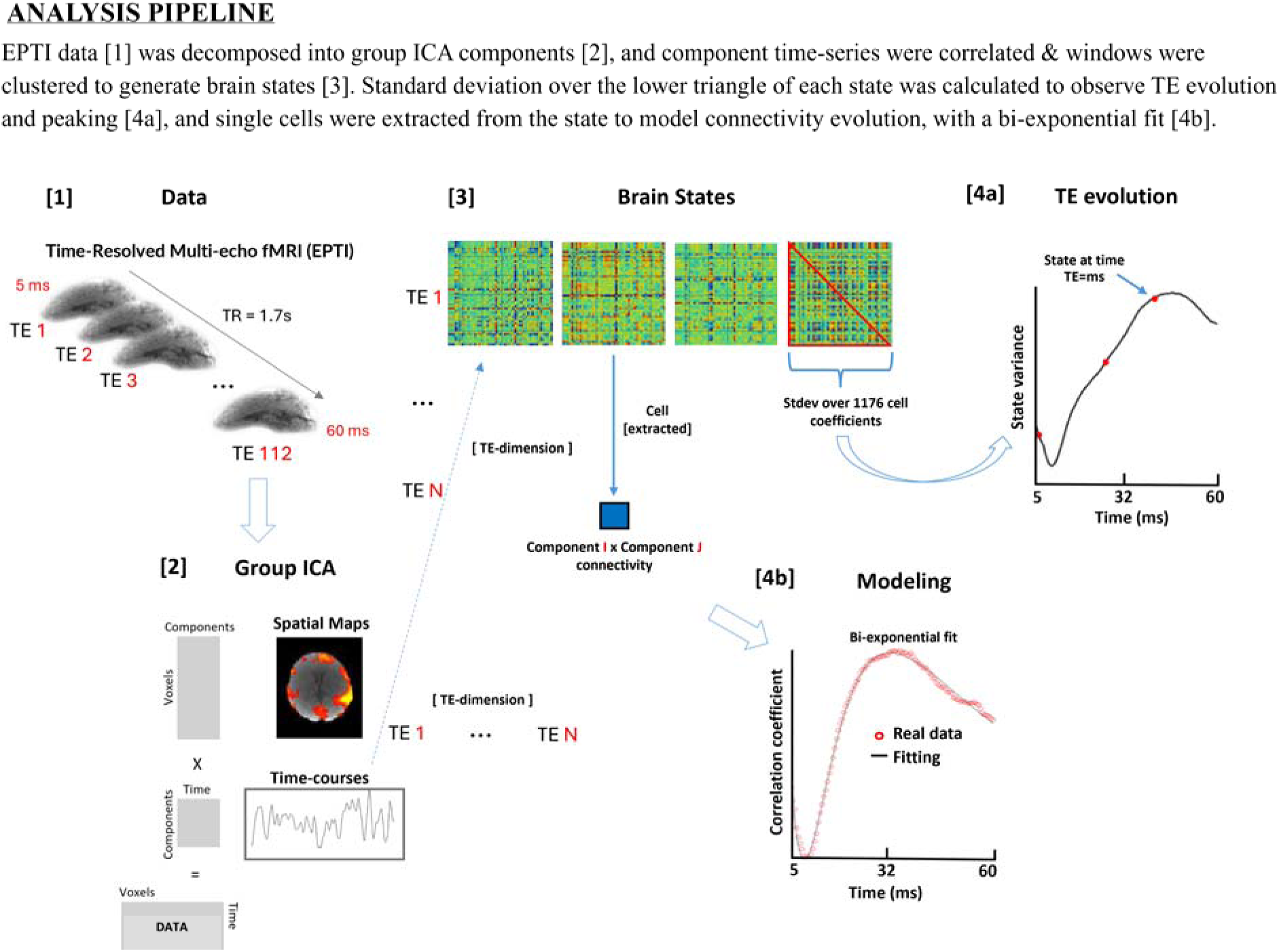

## 1 Introduction

Multi-echo functional magnetic resonance imaging (fMRI) collects multiple echo volumes within each repetition time (TR). Averaging these echo volumes after weighting each echo volume by T2* allows enhancement of the volume signal-to-noise ratio (SNR), providing a buffer against signal dropout from accelerated T2* signal decay in brain regions proximal to air-tissue cavities (Posse, 1999; Poser, 2006; Olman, 2009). Multi-echo volumes also provide TE-dependent information that can be leveraged for more advanced characterization. Recently developed advanced multi-echo imaging sequences, Echo Planar Time-resolved Imaging or EPTI (Wang, 2019; Dong, 2024), allow acquisition of over 100 echo volumes within a single timepoint (i.e., repetition time (TR)), which enables modeling of ‘temporally-resolved’ signal on a sub-millisecond time scale across TEs. Studies show that signal intensity in brain regions susceptible to signal dropout is largely recovered in EPTI, due to the presence of short T2* signal components or proton-density-weighted (PD) information in the earliest TEs, with evolving contrasts to predominantly T2*-weighted information in longer TEs (Dong, 2024; Krishnamurthy, 2023).

For example, images at early echo times with minimal signal losses from T2* relaxation are regarded as PD-weighted (Symms, 2004; Tofts, 2003; Krishnamurthy, 2023), which can be quantified or obtained by extrapolating image signal intensity to TE=0 (S0) (Tofts, 2003). Other studies suggest that the resolved PD-related signal intensity reflects microstructural or cellular processes, including potential changes in water content (Stroman, 2002). This localized increase in signal intensity occurs primarily at short TEs, and in areas which normally would experience signal dropout or relaxation at longer TEs (Stroman, 2008; Figley, 2010; Ng, 2006). While commonly regarded as noise in T2*-weighted fMRI, PD information provides a quantitative measure of spin density, or relative percentage unit (pu) of water protons. Thus, it can provide a measure of structural integrity potentially related to neural-related signal changes (Stroman, 2002; Stroman, 2008). Structural properties that may be captured by proton-density involve tissue type differentiation, tissue atrophy in grey matter, de-myelination in white matter, or swelling in clinical cases (Kitzbichler, 2021; Tofts, 2003; Symms, 2004). Research suggests that the relaxation rates of water components at these early echo times (∼10-30ms) are more related to cellular water dynamics (Anastasiou & Hall, 2004; Chang, 1972; Tofts, 2003), than more freely-flowing water relaxation at longer TEs, that might be observed in BOLD or CSF. Here, we examine how connectivity-derived brain states evolve across echo times, without removing early-TE signal contributions.

In conventional multi-echo fMRI, the BOLD signal (ΔS) increases with TE until reaching a maximum at TE=T2*, while the non-BOLD signal decreases monotonically with increasing TE. A diverging relative signal difference from a baseline mean (ΔS(TE_n_) = S(TE_n_)_activation_ – S(TE_n_)_baseline_) indicates a relative signal increase from neuronal-related task activation or resting-state neuronal activity (Kundu, 2012; DuPre, 2021). Parameters are estimated on a voxel-wise basis by fitting signal changes across TEs to a mono-exponential model: S(TE_n_) = S0 * exp(-TE_n_/T2*). The two parameters estimated from this fit include: 1) T2* decay rate in ms and 2) initial signal intensity approximated to TE=0 (S0) (Kundu, 2012; Bandettini, 1994; DuPre, 2021). As echo time is directly related to this relative increase (ΔS) in BOLD contrast amplitude, longer echo times observe higher peak BOLD signal, compared to shorter echo times with less BOLD (Boursianis, 2021; Menon, 1993, Kundu, 2012). Although short TEs are historically susceptible to noise artifacts (e.g., in-flow effects, physiological noise, and head motion) (Glover, 1996; Bright & Murphy, 2013; Kruger & Glover, 2001), short TEs may still contain valuable signal information. The slow-growing BOLD signal may be observed in mid-longer TEs (Miller, 2001; Ogawa, 1990; Bandettini, 1994); whereas, short TEs may contain different signal information. While multi-echo volumes are typically combined to resolve signal decay, very few studies have explicitly examined the evolution of functional signal information across TEs, on a network-level (Yuan, 2021; Van de Moortele, 2008; Zhao, 2024; Feng, 2025).

Short echo times (∼11-15ms) also contain early signal information that precedes the BOLD response, such as signals related to tissue microstructure and cellular properties (Stroman, 2002; Stroman, 2008; Figley, 2010; Krishnamurthy, 2023; Dong, 2024). T2* relaxation rates of these microstructural signals tend to be faster and captured at these shorter TEs (Stroman, 2002; Kijowski & Chaudhary, 2014; Du, 2013; Afsahi, 2021; Brix, 1990). As T2* decay can be sensitive to tissue type and cellular properties (Boursianis, 2021; Li, 2012), other studies have modeled T2* as a two-component sum of cellular tissue contributions and vascular sub-components (Ulrich & Yablonskiy, 2017; Im, 2025). Thus, bi-exponential models can be used to model relaxation time of short and long signal components with a non-rapid exchange rate (Stroman, 2002; Brix, 1990; Anastasiou & Hall, 2004).

In this study, we applied a higher-level implementation of a bi-exponential model to theoretically model short-TE signal connectivity that decays across TEs, and long-TE signal connectivity. Functional network connectivity (FNC) (a measure of neural-related BOLD signal correlation between two regional networks over time) (Tagliazucchi, 2012; Allan, 2015) has been validated by TE-dependence through covariance (Gonzalez-Castillo, 2026).

The decay rate term is deemed as FNC_decay_ (decay in connectivity by the short and long components) and FNC_TE=0_ (initial connectivity magnitude at TE=0). As a phenomenological model, we extend the model input from typical signal intensity (S(TE_n_)) changes across TEs used in conventional multi-echo models (Kundu, 2012). As conventional models explore relative changes in BOLD contrast on a voxel-wise level (ΔS(TE_n_)), we explore higher-level signal changes on the FNC level. In our pipeline, denoised voxels were mean-centered and linear-detrended prior to GICA, and our higher-level GICA components are also z-scored, to remove the time-series mean. Thus, when the mean = 0, the variance or standard deviation of the time-series holds the bulk of the signal information. At this stage, both the mean and variance of our GICA component time-series are independent (uncorrelated) across TEs.

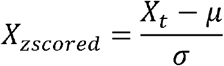

The standardized covariance between the z-scored GICA components are then computed via FNC:

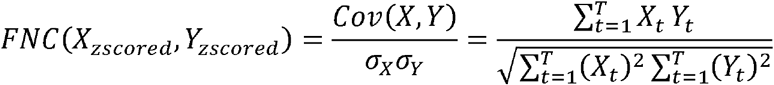

Theoretically, if the FNC between the two GICA components contains BOLD information, the connectivity would increase to peak around T2* (i.e., a constant representing the rate of signal decay over time). If the FNC between two GICA components holds non-BOLD or other information, the connectivity would decrease or show alternative trends (Gonzalez-Castillo, 2026).

Here we present the canonical mono-exponential decay model (S0 = initial signal intensity at TE=0, T2* = signal decay rate):

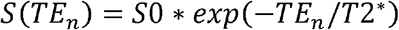

And in this study, we present a higher-level model for modeling FNC connectivity behavior, that builds off of the canonical mono-exponential signal decay (S0 = fnc_TE=0_, T2* = fnc_decay_), to avoid term confusion with canonical modeling:

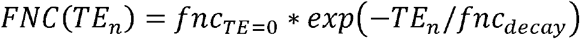

Per the characterization of PD versus BOLD-weighted information in T2*-weighted multi-echo data (Stroman, 2002; Dong, 2024; Krishnamurthy, 2023; Kundu, 2012), we assume that the short-TE connectivity is decaying (FNC(TE_n_)_NonBOLD_) and the long-TE connectivity (FNC(TE_n_)_BOLD_) is growing in influence across TEs. We model the increasing model by flipping the exponential decay model, along its y-axis (-exp).

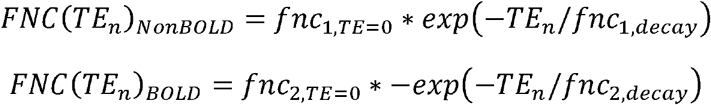

We then take the squared sum of the above two exponentials (decaying and increasing models), to model the combined signal evolution across TEs (FNC(TE_n_)_TOTAL_):

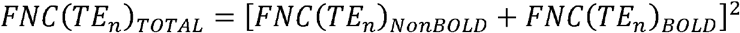

This higher-level phenomenological model was necessary to model TE dynamics as a result of FNC. At this stage, differentiating between non-BOLD and BOLD information is not dependent on a relative baseline signal mean (ΔS(TE_n_) = S(TE_n_)_activation_ – S(TE_n_)_baseline_), but rather on the covariance between two uncorrelated time-series (FNC(X_zscored_,Y_zscored_)) – where the time-series information dictates the BOLD versus non-BOLD relationship observed across TEs. For simplicity, we do <u>not</u> use these terms interchangeably throughout the manuscript (S0_1_ = fnc_1,TE=0_, S0_2_ = fnc_2,TE=0_, T2*_1_ = fnc_1,decay_, T2*_2_ = fnc_2,decay_), until the discussion section and modeling simulations for interpretation. It is important to note that these terms should not be interpreted as S0 or T2* in the canonical sense, but rather as parameter terms that model and separate signal properties responsible for alternative trends in exponential decay in FNC connectivity.

Despite extensive work on multi-echo denoising and signal characterization, little is known about how dynamic functional connectivity states evolve across echo times. In particular, it remains unclear whether TE-dependent signal composition alters the structure, variance, and temporal behavior of connectivity states. Here, we address this gap by treating echo time as an explicit dimension in dynamic state analysis.

## 2 Materials and Methods

### 2.1 Data acquisition and preprocessing

#### Acquisition parameters

Single-shot gradient-echo EPTI fMRI data was acquired on a 3T Siemens Prisma scanner, with a 32-channel head coil on six healthy volunteers (2 males, 4 females, ages 22 – 34 yrs old), with a consented institutionally approved protocol. A total of 112 simultaneously-acquired T2*-weighted echo volumes were collected, with TEs ranging from 5-60ms using a spatio-temporal controlled aliasing in parallel imaging (CAIPI) sampling trajectory to generate time-resolved distortion- and blurring-free echo volumes (Wang, 2019; Dong, 2024). Acquisition parameters included isotropic voxels with resolution 3×3×3mm^3^, slice thickness=3mm, FOV=216×216mm, number of slices=42, TR=1.7s, number of echoes=112, ESP=0.49ms, SMS=2.

#### Preprocessing

The preprocessing steps included EPTI data subspace reconstruction (Dong, 2020), slice-timing-correction, motion-correction (*3dvolreg, 3dAllineate*, AFNI) (Cox, 1996), volume alignment to T1, regression of physiological and MRI noise components on raw native-space data (*fsl_regfilt*, FSL) (Smith, 2004), warping to MNI space (*fsl_applywarp*, linear and nonlinear transformations, FSL), and smoothing to 8-mm FWHM (*3dBlurToFWHM*, ‘ACF’ smoothing option for Gaussian-exponential model, AFNI).

Transformations were first estimated from the optimally-combined echo volume and subsequently applied to each echo volume.

#### Denoising

MELODIC ICA (FSL, spatial-smoothing=8mm, prior to ICA) was applied to the raw, optimally-combined echo volume and 1^st^ echo volume (TE=5ms) for each participant in native-space. We then manually identified physiological (cardiac, respiratory, WM), susceptibility-motion, and MRI-related noise components from the optimally combined echo volume, and MRI noise components from the 1^st^ echo volume - since MRI noise components were noticeably cleaner in the first TE. Noise components were manually classified using Griffanti’s guide (2017) as a reference to identify fMRI artifacts through evaluation of the component spatial maps, time-courses, and power spectra. Finally, we generated a design matrix with signal and noise components and filtered out the noise components from each echo volume using partial regression (*fsl_regfilt*).

### 2.2 Group Independent Component Analysis (GICA)

All GICA and dFNC analyses were conducted using the GIFT toolbox (*GIFT, v4.0*). Prior to ICA, voxel-wise data was pre-processed by removal of the image mean per time-point and detrended. ICA-based decomposition of the preprocessed data resulted in 49 independent components (ICs) (a subset of Neuromark 1.0), via GIFT’s spatially-constrained ICA algorithm (sc-ICA). This algorithm utilizes a semi-blind fixed-point iteration algorithm to constrain each decomposed network component to regions defined by the group-validated Neuromark atlas (Du, 2020; Calhoun, 2001). As we excluded the cerebellar components (ICs 50 - 53) within the original Neuromark template, our 49 component networks were inclusive of 6 brain networks: subcortical (SC), auditory (AUD), sensorimotor (SM), visual (VIS), cognitive-control (CC), and default-mode (DM). Group ICA (GICA) was conducted across all 112 echo volumes, and GICA components were back-reconstructed to each individual echo volume, via spatial-temporal regression. Spatial-temporal regression takes the aggregate GICA spatial maps and regresses onto the actual data (i.e., individual echo times) to generate echo-specific time-courses. The second part of spatial-temporal regression regresses the aggregate GICA time-courses onto the data to generate echo-specific spatial maps. After back-reconstruction, the final echo-specific GICA component spatial maps and time-courses were scaled to z-scores. Importantly, echo times were treated as a structured dimension rather than independent samples, allowing preservation of cross-TE dependencies during decomposition.

### 2.3 Dynamic Functional Network Connectivity Analysis (dFNC)

Prior to dFNC, GICA component time-courses were detrended (any residual linear trend and mean removed), de-spiked, and low-pass-filtered with a ceiling of 0.28 Hz. A window size of 27 TRs was selected to balance temporal resolution and estimation stability, consistent with prior dFNC studies (Allen, 2014). Gaussian (alpha=3 TRs) windowing function was applied during dynamic FNC (dFNC). The Gaussian window iterated over component time-course pairs (with a step-size of 1 timepoint (TR)), resulting in a whole-brain matrix (NxN= 49×49) of network correlations (L1-regularized sliding window Pearson’s correlation (SWPC)) within each dFNC window. We performed L1 regularization (100 iterations) on the resulting dFNC matrices to enforce sparsity and reduce noise within the windowed correlation matrix.

**DIAGRAM 1.**
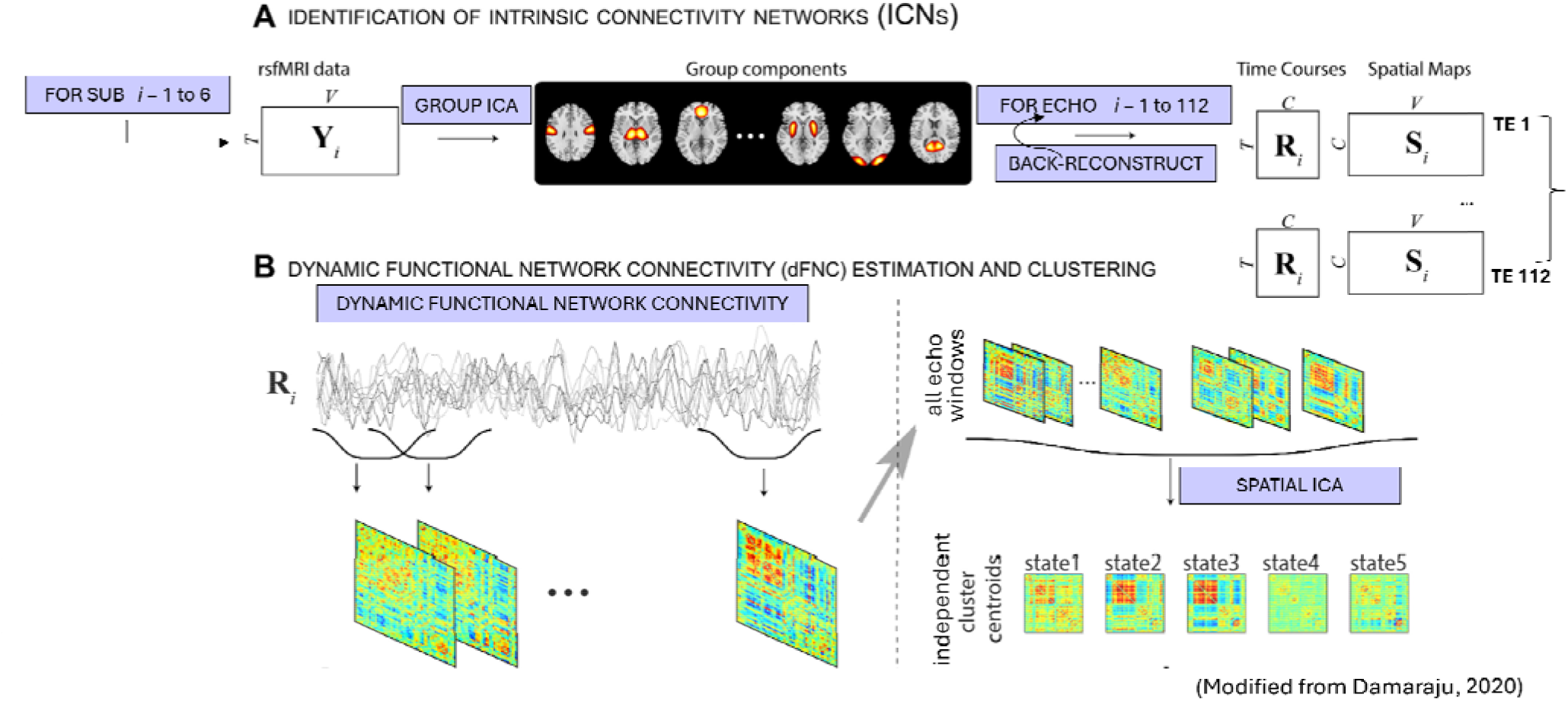
TE-dependent Group TCA and Dynamic Functional Network Connectivity Analysis

### 2.4 Spatial ICA State Clustering

#### Generating distinct brain states

The optimal number of clusters (*k*) was estimated using the elbow criterion with window exemplars (windows of maximal standard deviation) extracted from each echo time (*GIFT, v4.0*). DFNC windows were then concatenated across echo times (on a within-subject level) and across echo times and subjects (on a group-level). After concatenating the windows, second-order decomposition (i.e., 2D singular value decomposition (SVD)) was performed across the time dimension to generate a set of rank-1 matrices (i.e., distinct brain states), with the following dimensions: ((space x TEs) x k). Higher-order decomposition (i.e., independent component analysis (ICA)) was then performed on the decomposed brain state to generate a set of statistically independent brain states, that have ‘maximal’ spatial independence from each other. The ‘Extended Infomax’ algorithm (Lee, 1999) was used to perform ICA, as it generates sources (brain states) with higher CNR and is largely insensitive to random initialization (*GIFT, v4.0*, Iraji, 2020). This method of concatenating subject-specific windows prior to spatial ICA clustering (Soleimani, 2025), allowed us to preserve the TE dimension during spatial ICA clustering.

#### Generating time-courses (not shown)

Conducting ICA resulted in a set of final state sources with the ‘TE’ dimension preserved: ((k x TEs) x 1176 [lower-triangle coefficients]). In order to generate corresponding time-courses with each state source, the ‘weighted-influence’ that each state source (X_i_) has on each dFNC window across TEs (x_TEi_ x_TEn_) is evaluated. This weighted time-course can be generated either by: 1) multiplying the mixing matrix (‘A’) by the ‘lambda’ output (singular values) from SVD: e.g., (k x k) * (k x TEs [windows]) – to reconstruct the original time-courses, or 2) regressing the state sources onto each dFNC window to obtain a time-course of beta-coefficients that can then be thresholded to generate a state transition matrix, for each TE (Soleimani, 2025).

**DIAGRAM 2.**
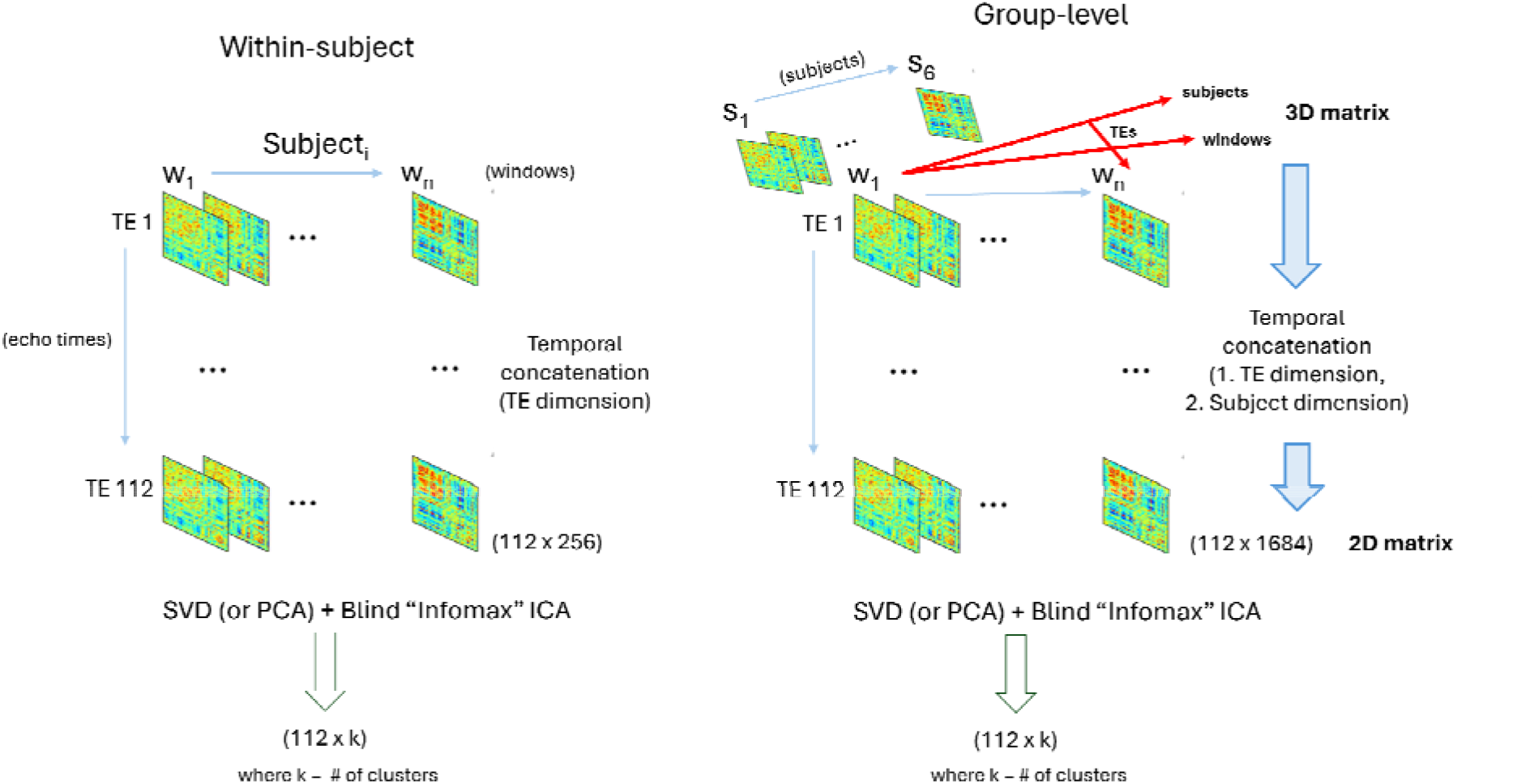
Spatial ICA State Clustering Analysis across TEs

### 2.5 A Note on Preprocessing

For multi-echo preprocessing, transformations (e.g., motion-correction, volume alignment, or warping) should be estimated from a chosen volume (such as, the bias-field-corrected optimally-combined echo volume) and the same transformation should be applied to individual echo volumes – to maintain integrity of signal information across TEs. Global signal and other signals related to the baseline mean intensity across voxels also may contain neuronal-related information, and thus should be evaluated more carefully before any global regression (Gonzalez-Castillo, 2026). In EPTI, the signal component time-series at early echo times contain more high-frequency bands than those at longer echo times. Thus, we minimally preprocessed our data and, for the GICA components, used a lower polynomial order (linear) for detrending and a higher threshold for the low-pass filter (0.28 Hz). In subsequent statistical analysis after preprocessing, we recommend leveraging the dependence of the echo volumes, rather than treating each individual echo volume as independent (e.g., concatenating the TEs during SVD/PCA, group ICA, or any iterative algorithm) – to avoid introducing additional noise across the TE dimension. If for some reason, you want to preserve correlated variance across TEs (e.g., during voxel-wise preprocessing or denoising), we recommend not scaling by the standard deviation.

### 2.6 Statistical Analysis

#### Within-subject state analysis

Within-subject brain states were generated from conducting spatial ICA on the time-concatenated dFNC windows across all echoes belonging to a participant. Concatenated states across TEs were then min-max normalized to convert state ranges from original ICA units to a more interpretable range of –1 to 1, without altering the covariance across TEs. State variance was analyzed across TEs by calculating the standard deviation across all 1176 (vectorized lower-triangle) state coefficients at each TE. The variance time-course was then smoothed with a moving average over TEs. The peak TE was calculated as the echo time with the maximal value of the variance time-course across TEs. State differences were calculated across the echo time range, by calculating the sum of squared differences across state coefficients between the peak TE (maximal variance) and each state at echo time t_i_.

#### Group-level state analysis

Group-level states were obtained from concatenating dFNC windows from all echoes and subjects across the time dimension and conducting spatial ICA on these time-concatenated windows. Group-state exemplars were calculated by 1) calculating the Euclidean distance between each state (t_n-1_ to t_n_) across time (TEs), and 2) taking the absolute derivative (difference) of the Euclidean distance time-series, to obtain clear peaks at certain TEs – where the state information changes significantly from prior TEs.

From this matrix of group-state exemplars (∼30 exemplars per group state), 3 exemplars were chosen from the beginning, middle, and end of the exemplar matrix. The a-priori chosen exemplar TEs were then mapped back to their state variance (standard deviation) time-course, which was also smoothed over TEs. Because these exemplars were chosen independently of the state variance time-course, they do not necessarily correspond to the exact location of peaks or troughs in the variance time-course – rather representing points in time where the state information may change the most.

### 2.7 Model Simulations and Fitting to Real Data

To evaluate the underlying mechanisms behind the unique behavior of evolving trends in dynamic connectivity across echo times, we implemented three model simulations using the bi-exponential model (see Introduction). Our modification to Stroman’s bi-exponential model (2002), takes the function a bit further and calculates the squared sum of these two exponentials to model dFNC signal connectivity across TEs.

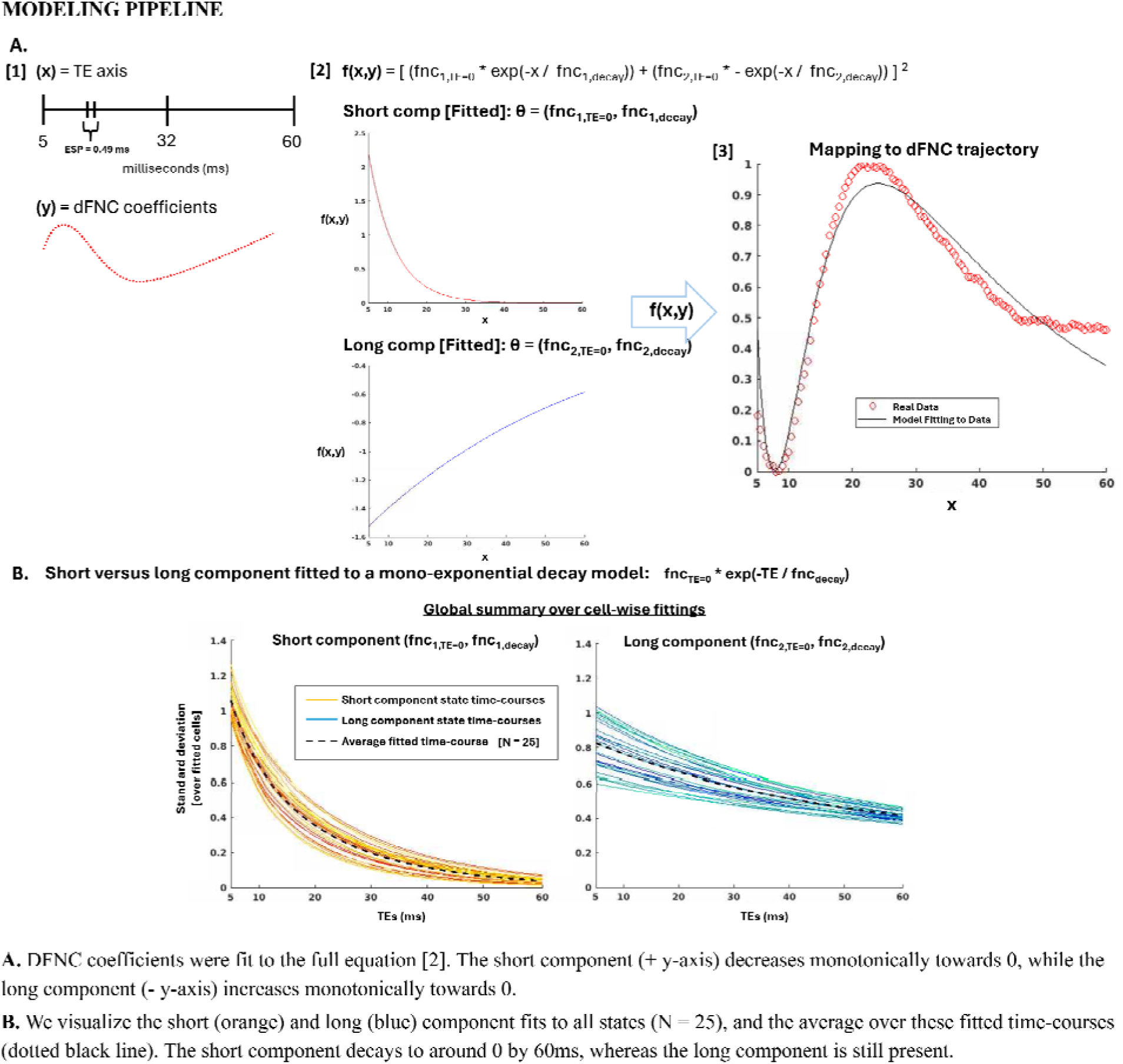

#### Model simulations

For the model simulations, we generated 3 models with different assumptions about the initial signal intensities of the two signal components at TE=0: 1) Model 1 assumes that S0_1_ > S0_2_, 2) Model 2 assumes that S0_1_ < S0_2_, and 3) Model 3 assumes that S0_1_ = S0_2_. All models assume that T2*_1_ decay is shorter than T2*_2_ decay. For simplicity, we keep the T2*_1_ decay at 10ms and only modulate the T2*_2_ decay rate from 10ms – 60ms, during simulations. We decided to use this ms range as an example for the simulations – to explore how differences between the short and long component decay rate may influence the magnitude of the simulated signal response.

#### Model fitting to real data

We extend the model to matrix cell-wise functional connectivity coefficient time-courses across TEs. These cells are extracted from the resulting brain state matrices, from spatial ICA of dFNC. To calculate a relative difference in FNC from a more conventional TE (i.e., FNC at TE=32ms), we scaled the dFNC time-course as following: FNC_relative_(TE_n_) = (FNC_i_ – FNC_TE=32ms_) / FNC_TE=32ms_. We then min-max normalized the resulting scaled dFNC time-course, to a range of 0 – 1. During nonlinear least squares fitting (*lsqcurvefit*, ‘levenberg-marquardt’ solver algorithm [MATLAB], echo times (ms) = x (independent data), normalized dFNC time-course = y (observed data)), we explicitly fit each cell-wise dFNC time-course to the 3 models defined during simulations. Model 1 initial parameter estimates (θ_0_) were (θ_1_ = [S0_1_ = 300, S0_2_ = 100, T2*_1_ = 10ms, T2*_2_ = 60ms]), model 2 initial parameter estimates were (θ_2_ = [S0_1_ = 100, S0_2_ = 300, T2*_1_ = 10ms, T2*_2_ = 60ms]), and model 3 initial parameter estimates were (θ_3_ = [S0_1_ = 100, S0_2_ = 100, T2*_1_ = 10ms, T2*_2_ = 60ms]). The dFNC time-course was fit to the squared sum of the bi-exponential model function (see Introduction) and a set of optimal predicted parameters (θ_pred_= S0_1_, S0_2_, T2*_1_, T2*_2_) was estimated for each model. As nonlinear least-squares fitting minimizes the error between the actual data and the predicted data, we chose the model predictions that produced the lowest residual error norm ( ∑ (y_pred_ - y_actual_)^2^).

Fitting the data to the three model initial parameter estimates (θ_1_, θ_2_, θ_3_) reduced the need to search for optimal initial parameters, and allowed the model fit with the lowest residual error norm to be chosen. To exclude bad fittings and non-convergent iterations from ‘successful’ iterations, we excluded cell-wise fittings that: 1) did not converge, 2) predicted a long T2*_2_ decay value that exceeded the length of the scan, and 3) predicted parameter estimates that were negative. Finally, a set of model performance statistics were calculated for all cell-wise fittings: goodness of model fit (R2 = 1 – (SS_residual_ / SS_total_), where SS = sum of squares), residual error norm, total number of function evaluations needed to fit the model, and total number of ‘successful’ iterations (converged cell-wise fittings / total cell-wise fittings, in which total cell-wise fittings = converged + non-convergent or excluded). There were no lower or upper bounds for any of the parameters included in the model fittings, since including parameter bounds tended to restrict parameter estimates and negatively affect accuracy (i.e., res. error).

## 3 Results

### 3.1 Within-subject brain state analysis

Within-subject brain states were visualized at TE’s of 5ms (TE_1_ or first TE), 32ms (TE_56_ or middle TE), and 60ms (TE_112_ or last TE). The max centroid was calculated as the state with the maximal standard deviation at any particular TE. We observed that states at short TEs (i.e., < 32ms) tended to have diverging whole-brain correlations, compared to states observed from ∼32 – 60ms. The max centroid (peak of state variance) showed similar state organization to early 5ms states (Fig. 1, part A), states at 32ms (conventional range) (Fig. 1, part B), and states at 60ms (longest recorded TE) (Fig. 1, part C).

**Figure 1.**
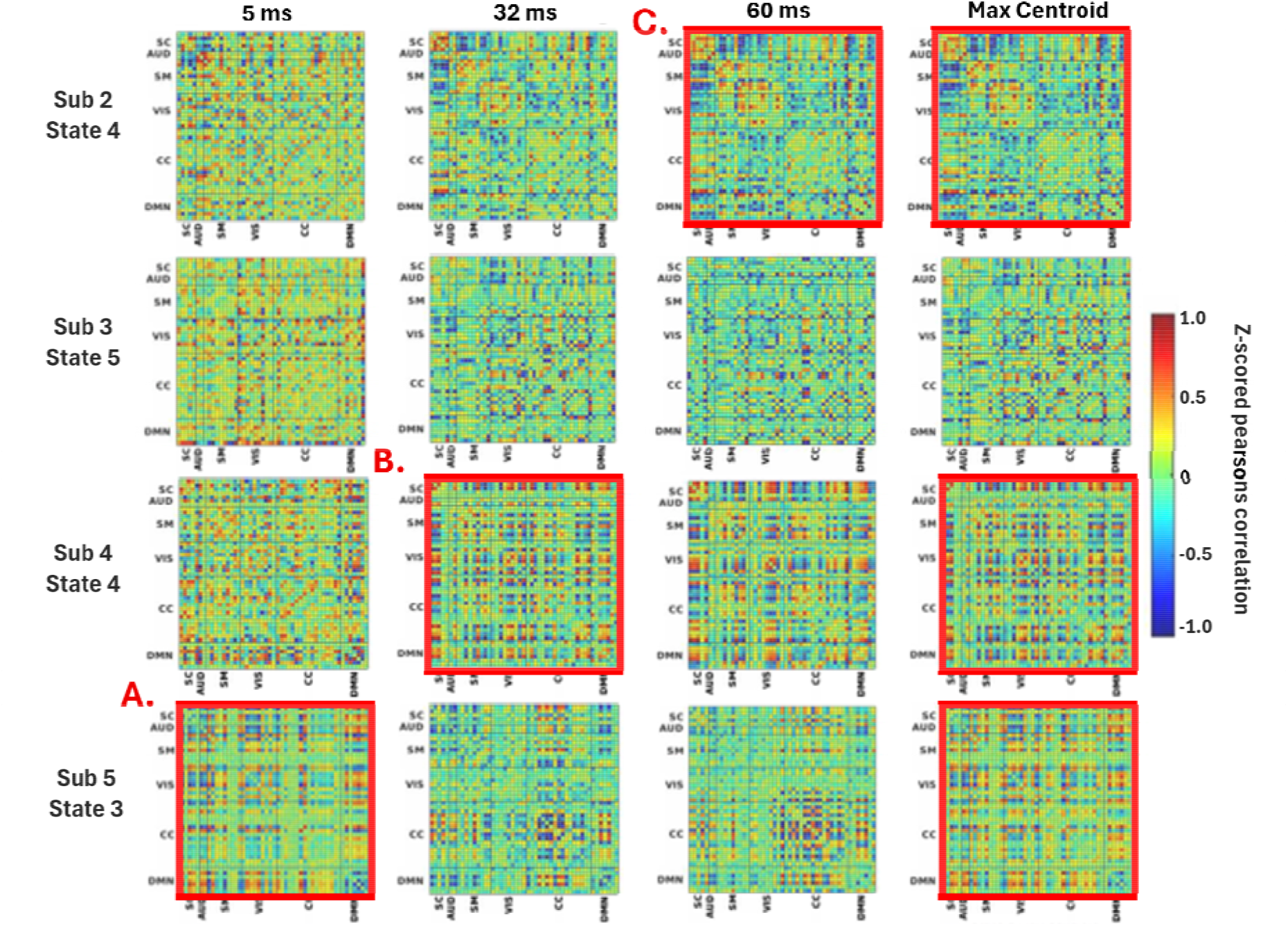
Visualizing within-subject states and peak TE state centroids. Max centroids appeared similar to states at 5ms, 32ms, and 60ms - for TEs at the beginning, middle, and end of the acquired TE range. The max centroid represents the slate alt ‘peak TE’ or maximal stale variance.

Despite their divergent appearance, these early states still contained some resemblance of functional organization. Given the large influence of PD-weighting at shorter TEs, these early states may reflect distinct signal contributions observable at short echo times for the first time.

State variance time-courses for each subject and their corresponding states were visualized below (Fig. 2, part A). Different states tended to show distinct changes in state variance, with peaking behavior at different echo times. States that peaked at conventional TEs gradually increased in variance over time, reaching a plateau in variance (i.e., peak TE) with a slight decrease in variance after the peak [1A]. Early-TE states peaked in variance at the earliest TE and then decreased over time [2A].

**FIGURE 2.**
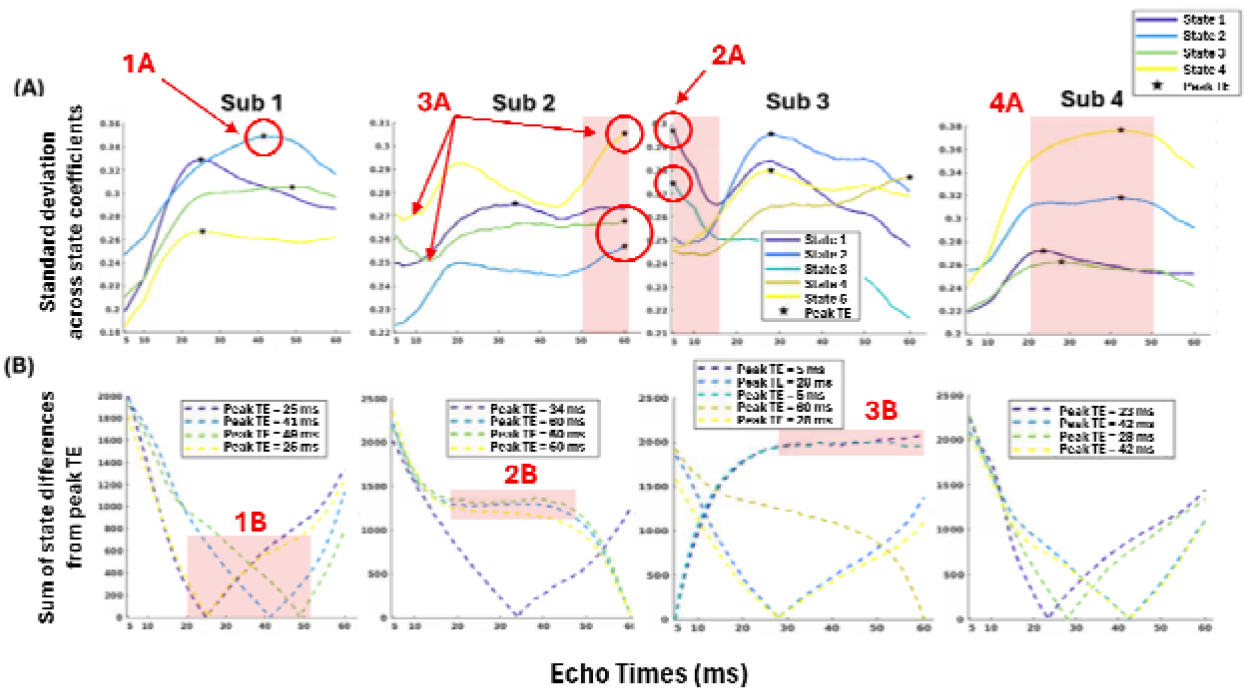
Evolution of subject-level slate variance and state differences across echo times. Fig. 2A. Slate variance lime-courses and peak TEs of maximal variance (black dots) were visualized. Fig. 2B. Stare differences from the peak TE {x-intercept or y-0) were also visualized.

Late-TE states dipped in variance after this early TE and gradually increased in variance over time – reaching peak at the latest echo time [3A]. States that peaked in conventional TE range (e.g., ∼20 – 50ms) tended to show more simple concave, curved trajectory with a clear, single peak [4A], while states that peaked at unconventional early (e.g., 5 – 15ms) or late TEs (e.g., 50 – 60ms) tended to show more complex trajectories in variance with local peaks and dips in variance along the echo time course [2A, 3A]. States peaking at early or late TEs are states that are less likely to have been detected in a conventional multi-echo sequence with TEs ranging from ∼20-50ms.

These complex changes in variance are possibly a result of evolving signal properties across TEs.

All state differences diminished rapidly to 0 at the peak TE (x-intercept at y=0) (Fig. 2, part B). Simple trajectories with peaks in the conventional TE range showed convex dips surrounding the peak TE [1B]. States that peaked at 60ms showed a cubic trajectory with state differences that plateaued in the middle TE range and diminished quickly to 0, as approaching the peak TE [2B]. States that peaked at 5ms, showed steep increase in state differences with a clear plateau across longer echo times [3B]. Many states belonging to a single subject also shared similar trajectories in state differences, and similar peaking times and behavior. State differences changed rapidly around the peak TE; however, for states that peaked at early or late TEs, plateaus in variance in the middle TE range suggested inherent similarity of state variance around the conventional TE range.

### 3.2 Group-level brain state analysis

We visualized the state variance time-course from all group states (N = 4), and independently mapped a-priori exemplar TEs to the state variance time-course (Fig. 3). The exemplars (red dots) represent points in time where the state information changes the most from prior TEs. Notably, these exemplars (despite independence from the state variance time-course) corresponded to TEs near the beginning of the acquisition, close to peaks or troughs in variance, and directly along slopes of variance change or along plateaus in variance (Fig. 3). In the pre-hoc (a-priori) analysis, exemplar TEs (i.e., ∼6ms, 26ms, 40ms) were also similar across group states, suggesting that certain TEs may be more prone to exemplar behavior than others.

**FIGURE 3.**
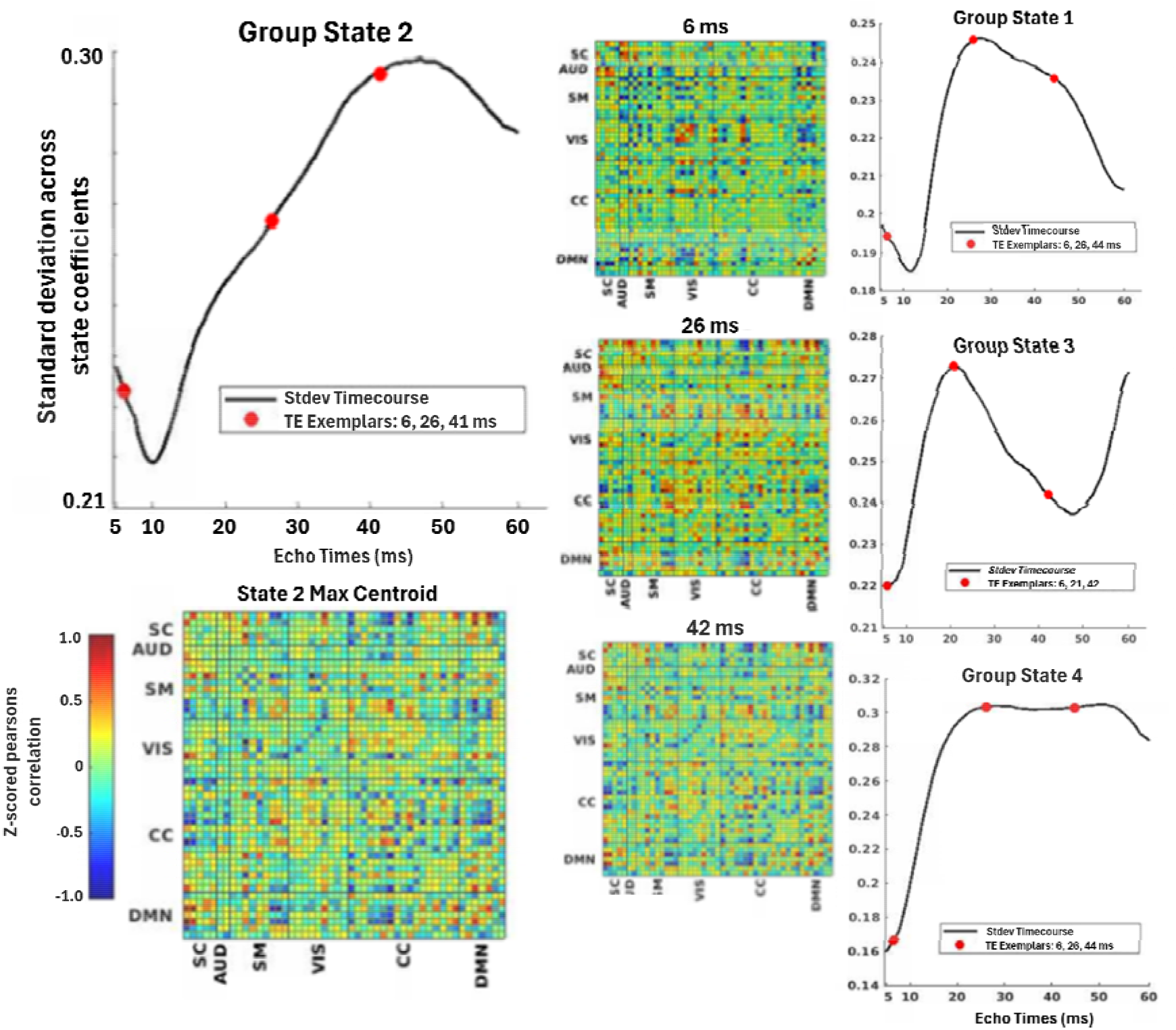
Mapping group-level state exemplars along the variance time-course. Group-level state variance time-courses and a set of group state exemplars (red dote;) were extracted for each state - revealing unique trends and similar points of transition.

In a post-hoc analysis (Supplementary Information, Figure 8), we performed PCA decomposition on the extracted states across the TE dimension and extracted time-courses for each of the PCA-decomposed states. This analysis more directly mapped real transitions in state across TEs to evolution of state variance. The state time-courses were then standardized and thresholded by maximum value across TEs, in order to generate the state transition vector. The first derivative (*diff*, MATLAB) of this state transition vector allowed direct mapping of state transition points onto the state variance time-course.

### 3.3 Modeling of dynamic FNC across echo times

In our model simulations, we found model 1 (S0_1_ > S0_2_) revealed an initial dip in connectivity at short TEs (Fig. 4A). Model 2 (S0_1_ < S0_2_) revealed an early bump in the total signal intensity with a gradual decrease across echo times (Fig. 4B). Model 3 (S0_1_ = S0_2_) revealed a bell-shaped curve, with signal peak at more conventional TEs in the middle-TE range (Fig. 4C). Alternative trends indicate the strong influence of the short component’s initial signal intensity (S0_1_), relative to the long component’s. Again, note that the use of two S0 and T2* terms and modeling the squared sum of a decreasing and increasing exponential should only be interpreted at the higher-level – when BOLD-related signal information has already been differentiated from non-BOLD.

**Fig. 4.**
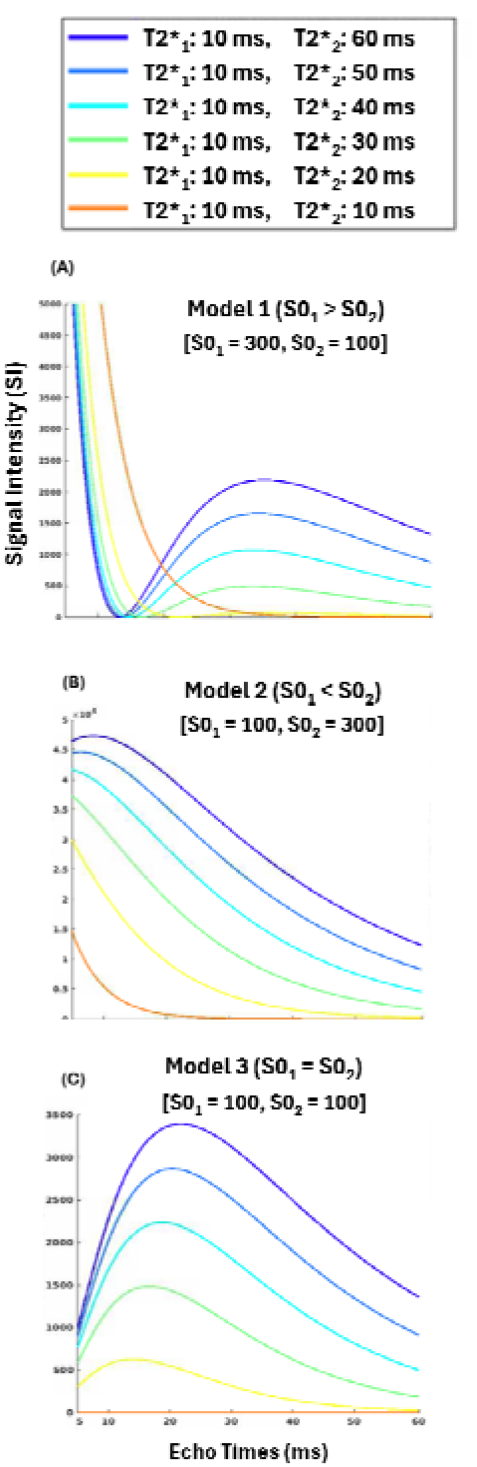
Three different model simulation scenarios 1-3 assume initial signal intensity al TE=-0 (arbitrary units) for the short cmnponc1n versus the long component. Short component decay (T2* 1) was kept constant at 10ms, while only the long component decay (T2*_2_ was modulated from I ll-61lms.

Our simulations suggest that relative differences in initial signal intensity of the short and long signal components (S0_2_/S0_1_) were responsible for the unique shapes in signal trends that we observed across echo times (e.g., initial dip, initial bump, or bell-shaped curve) (Fig. 4). The simulations also suggest that a substantial difference in T2* relaxation rate (i.e., between T2*_1_ and T2*_2_) was necessary to model these more ‘complex’ shapes in signal evolution. When the short and long components dFNC decayed at the same rate (e.g., T2*_1_ = T2*_2_ = 10ms, orange line in Fig. 4), a single exponential decay (or flat line) was modeled – suggesting that only a single component model is detected when components are exchanging at the same rate.

From the cell-wise fittings to the within-subject brain states, we visualized a single cell (network_i_-by-network_j_ dynamic connectivity) over TEs from all 6 subjects. As you can see, the trends observed in the model simulations were similar to those observed in our experimental data (Fig. 5): the bell-shaped curve when the initial FNC magnitude ratio (fnc_2,TE=0_/fnc_1,TE=0_) is close to 1.0 (Sub. 1), initial dip in connectivity when the long initial FNC magnitude is relatively smaller (fnc_2,TE=0_/fnc_1,TE=0_ < 1.0) (Subs. 2,4), and short-TE bump in connectivity when long initial FNC magnitude is larger (fnc_2,TE=0_/fnc_1,TE=0_ > 1.0) (Sub. 3). A two-component signal system can lead to delays in the decay rate (i.e., longer signal decay is reported) of the total signal due to the presence of the non-BOLD component (Van de Moortele, 2008). Thus, many of our long FNC decay (fnc_2,decay_) values were greater than 60ms – when the short component has a particularly strong influence (fnc_2,TE=0_ / fnc_1,TE=0_ ≈ 0.3 0.4).

**FIGURE 5.**
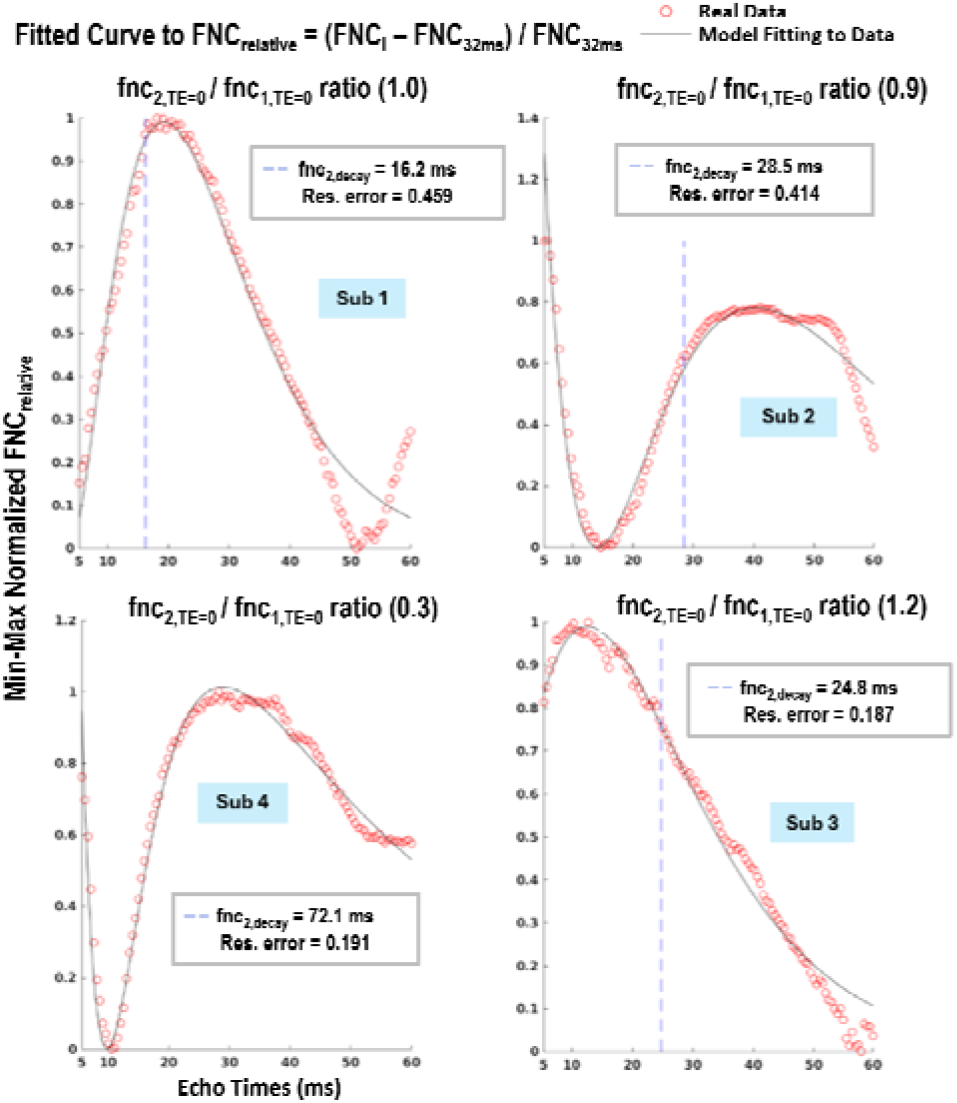
Model Fitting to cell-wise dFNC coefficient trajectories across TEs. Select lime-courses (red dots) and their-model fillings (black line) were visualize.cl. Each time-course represents the dynamic connectivity between two networks (cell-wise) evolving across echo times.

For a comprehensive evaluation of the model performance on real data, we conducted fittings on all 1176 coefficients per within-subject state. We then evaluated model performance across all states for a particular subject - as well as average model performance across all subjects (Table 1).

**Table 1.** Distribution statistics from model fittings 10 subject-level states. The performance statistics were generated along with cell-wise model fittings for all within-subject states, and gathered into a distribution. Statistics were included for all subjects, as well as the average over the statistic distribution medians (last line).

| Subjects | Model Fit (R2) | Residual Error | # of Function Evaluations | Converged / Total Iterations |
| --- | --- | --- | --- | --- |
| 1 | 0.89 (0.63 0.96) | 0.43 (0.13 1.16) | 88 (64 126) | 0.87 |
| 2 | 0.84 (0.53 – 0.94) | 0.56 (0.18 – 1.40) | 90 (65 – 130) | 0.85 |
| 3 | 0.89 (0.64 0.96) | 0.44 (0.14 1.16) | 89 (65 132) | 0.86 |
| 4 | 0.84 (0.48 0.94) | 0.57 (0.16 1.45) | 85 (60 125) | 0.84 |
| 5 | 0.88 (0.57 – 0.96) | 0.43 (0.12 – 1.15) | 85 (62 – 118) | 0.85 |
| 6 | 0.84 (0.62 0.93) | 0.87 (0.33 1.93) | 85 (65 122) | 0.93 |
| Average | 0.86 | 0.55 | 87 | 0.86 |

Reporting format: Median (Q1 – Q3)

Each cell in Table 1 provides the median (as a measure of central tendency) and first quartile (Q1) and third quartile (Q3) (as measures of deviation). The model performed sufficiently well, given the average statistics (bottom line): R2 fit = 0.86, residual error = 0.55, average number of function evaluations needed to converge = 87, and a success rate (converged iterations / total iterations) = 86%. The median model fitting statistics also did not vary greatly across subjects, revealing stability and generalizable performance across different states and subjects (Table 1).

We also evaluated state properties as a function of the estimated parameters from the bi-exponential model decay fitting. We evaluated the following state properties: 1) relative initial connectivity magnitude (fnc_2,TE=0_/fnc_1,TE=0_), 2) relative FNC decay rate (fnc_2,decay_/fnc_1,decay_), and 3) short FNC connectivity magnitude relative to long FNC decay rate (fnc_1,TE=0_/fnc_2,decay_). We included the last term (fnc_1,TE=0_/fnc_2,decay_ or S0_1_/T2*_2_) to assess properties from Stroman’s 2002 paper, which suggests that the short component’s signal intensity (S0_1_) may be relevant to microstructural signal properties represented by intensity (PD or extravascular water protons); whereas, BOLD is defined by the T2*_2_ relaxation term (Stroman, 2002). In order to evaluate these relative terms, we first calculated the ratio of the two terms (fnc_2,TE=0_/fnc_1,TE=0_), applied a log10 transform to the distribution of ratios, and min-max normalized to a range of 0 – 1. The resulting distribution statistics thus represent the normalized log ratio (i.e., log10(A/B) = log(A) – log(B)) or relative log difference. These state property distributions can be visualized in (Supplementary Information, Fig. 6).

In Table 2, we observed a significant inverse correlation (*r*(24) = -0.43, *p* = 0.03, 95% CI [-0.70, -0.04]) between relative initial FNC magnitude (fnc_2,TE=0_/fnc_1,TE=0_) and short initial FNC magnitude relative to long FNC decay (fnc_1,TE=0_/fnc_2,decay_). This indicates that three bi-exponential parameter terms (fnc_1,TE=0_, fnc_2,TE=0_, and fnc_2,decay_) may play a complex role in determining state connectivity dynamics.

**TABLE 2.** Pairwise pearsons correlation between relative ratio state parameters. Pairwise pearsons correlation between all stales and subjec.ls were conducted across the stale distribution medians **(N** – I – 24 d.o.f).

| *Relative Ratio of State Parameters* | $(\text{fnc}_{2,\text{TE}=0} / \text{fnc}_{1,\text{TE}=0}) \times (\text{fnc}_{2,\text{decay}} / \text{fnc}_{1,\text{decay}})$ | $(\text{fnc}_{2,\text{TE}=0} / \text{fnc}_{1,\text{TE}=0}) \times (\text{fnc}_{1,\text{TE}=0} / \text{fnc}_{2,\text{decay}})$ | $(\text{fnc}_{2,\text{decay}} / \text{fnc}_{1,\text{decay}}) \times (\text{fnc}_{1,\text{TE}=0} / \text{fnc}_{2,\text{decay}})$ |
| --- | --- | --- | --- |
| *R-coefficient: r(24)* | -0.20 | -0.43* | -0.16 |
| *p-value* | 0.33 | 0.03* | 0.46 |
| *95% CI [0.05,0.95]* | [-0.55, 0.21] | [-0.70, -0.04]* | [-0.52, 0.26] |

Significance: \* $p < 0.05$

## 4 Discussion

Given the small sample size (N=6), this study was intended as a methodological and exploratory analysis rather than a population-level inference. Our analyses reveal that echo times provide enhanced sensitivity allowing tracking of dynamic changes in brain state connectivity, variance, and other state properties.

Furthermore, we introduce a methodological approach of TE-concatenation that can be utilized to examine dynamic changes in brain state over TEs. Bi-exponential modeling of TE-dependent changes in connectivity confirmed a two-component signal system at the higher-level, and parameter evaluation suggests other properties within the signal (Supplementary Information, Figure 7). In a post-hoc analysis, regression of parameter maps (fnc_1,TE=0_, fnc_2,TE=0_, fnc_1,decay_, and fnc_2,decay_) onto TE-dependent states (N = 25 states) across echo times revealed unique temporal profiles per parameter term (Supplemental Information, Figure 7). And a secondary post-hoc analysis, evaluated parameter-dependent contributions to state connectivity dynamics, via state distributions (Supplementary Information, Figure 6).

The present study does not directly measure underlying biological processes. Rather, it demonstrates that dynamic connectivity states exhibit systematic and reproducible variation across echo times, and that a two-component model provides a useful framework for describing these patterns. Because signal changes are a function of both non-BOLD biophysical factors (e.g., water proton spin density, blood volume fraction, deoxyhemoglobin concentration) and BOLD (Kim & Ogawa, 2012; Blockley, 2013; Im, 2025), we do not consider all non-BOLD factors as noise in this study. Although we used this phenomenological model on higher-level processed dFNC state cells, the difference between the two signals evolving across TEs may be more accentuated by the processing induced by GICA and dFNC algorithms. Thus, voxel-wise modeling will require a different model – to properly separate bi-exponential signal decay. Structural and functional connectivity are inter-related (Santucci, 2025; Wang, 2022); however, we do not explicitly separate them within this study.

Per empirical literature, we present a few suppositions for interpretation of parameter terms. S0 terms (fnc_TE=0_, in this study) are related to signal intensity, and are inherently structural. T2* terms (fnc_decay_) are related to signal decay from sensitivity to contrast, and are inherently functional. We speculate that S0_1_ and T2*_2_ terms may be more closely related to PD and BOLD information (Stroman, 2002; Krishnamurthy, 2023). T2*_1_ term may be signal decay related to PD or cell water information (Tofts, 2003; Chang, 1972); while, S0_2_ term may be signal intensity related to BOLD information (Blockley, 2013; Havlicek, 2017). Again, these are only suppositions given the current literature, and would need to be quantified to hold any biological basis, in this context.

EPTI eliminates image distortion and corrects for dynamically changing B0-field inhomogeneities, improving robustness to motion and physiological noise (Wang, 2019; Dong, 2024). While long TRs (e.g., 1700ms) may minimize T1 effects and acquisition at short TEs (< 20ms) may minimize T2 effects, as is necessary to capture PD-weighted images (Tofts, 2003), PD-related information is not typically evaluated in a functional context with a time-series or across TEs. There is also still the possibility of residual T1, inflow, and physiological noise or motion that might persist after denoising. More advanced denoising pipelines (DuPre, 2021), or noise modeling may be used to remove these signal sources. Future pipelines may also seek to explicitly remove the PD component. Future studies should seek to employ more direct quantification of the signal across TEs, through biophysical modeling or direct acquisition of neural-related signals (e.g., EEG, CBF, CBV, CMRO2).

Ultimately, this study provides valuable methodology and insights for TE-aware state analysis, that can be directly applied at the basic science level, for a more informed interpretation of how functional connectivity evolves across echo times. A deeper look into signal decay parameter terms also provided a way to track how FNC evolution influenced state-specific properties. In the future, we hope TE-aware analyses can be implemented in translational studies to improve clinical interpretation and rehabilitation development.

## Author Contributions

M.H. preprocessing, preliminary and formal analysis, conceptualization, writing – first and final draft. A.B. preliminary and formal analysis and supervision – review and editing. B.B. conceptualization and supervision. S.W. methodology and conceptualization. Z.D. & F.W. data acquisition and MRI resources – review and editing. L.K. conceptualization, methodology, and supervision – review and editing. V.D.C. conceptualization, methodology, and supervision – review and editing.

## Acknowledgements

This material is based upon work supported by the National Science Foundation Graduate Research Fellowship Program under Grant No. (DGE-1937956). Any opinions, findings, and conclusions or recommendations expressed in this material are those of the author(s) and do not necessarily reflect the views of the National Science Foundation.

This work is also supported by the National Science Foundation through the CREST Center for Dynamic Multiscale and Multimodal Brain Mapping Over the Lifespan [D-MAP] (Grant No. 2112455). The MRI resources are supported by the NIH National Institute of Neurological Disorders and Stroke (Grant No. U24NS129893) and MGH Athinoula A. Martinos Center for Biomedical Imaging.

## Ethics Statement

Data acquisition involving human subjects was approved by MGB’s Partners Human Research Committee / IRB, and all participants signed informed consent, per MGB’s institutional protocol.

## Data Availability Statement

The data that support the findings of this study are available with permission from authors Z.D. and F.W., upon reasonable request.

