## Supplementary figures and images for "TE-aware state analysis and evolution in dynamic network connectivity"

### Supplementary Information

**Supplementary Information**

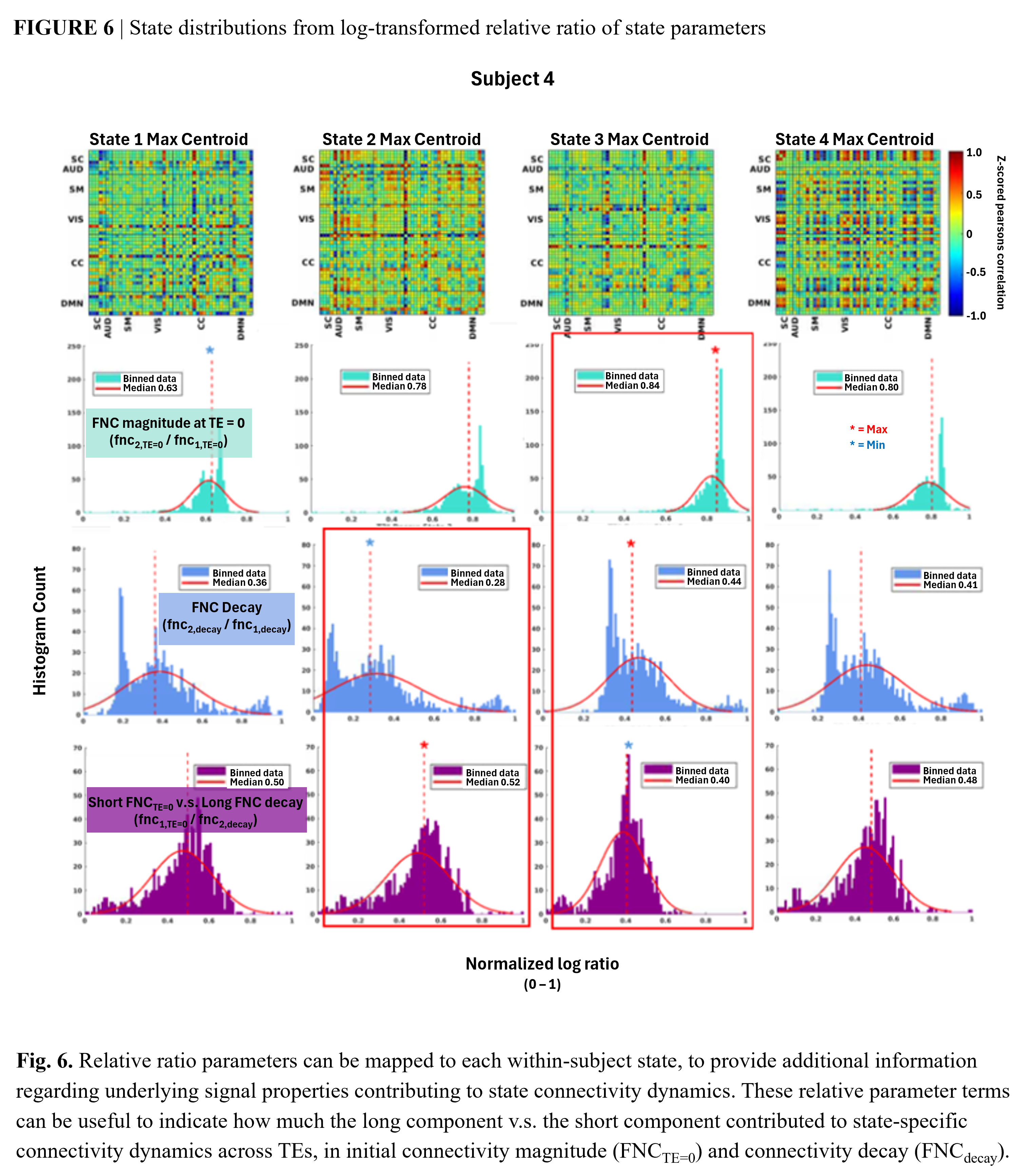

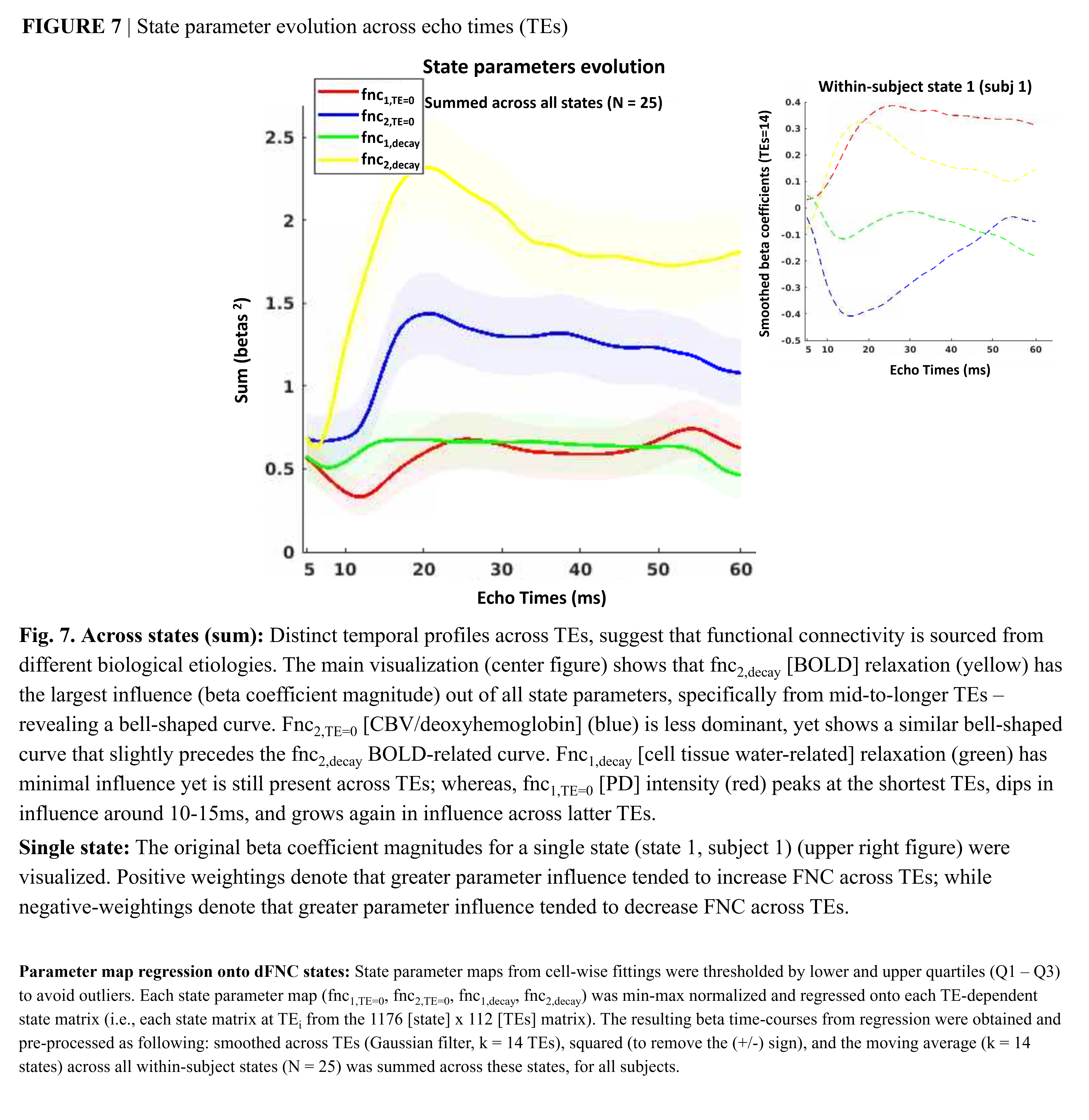

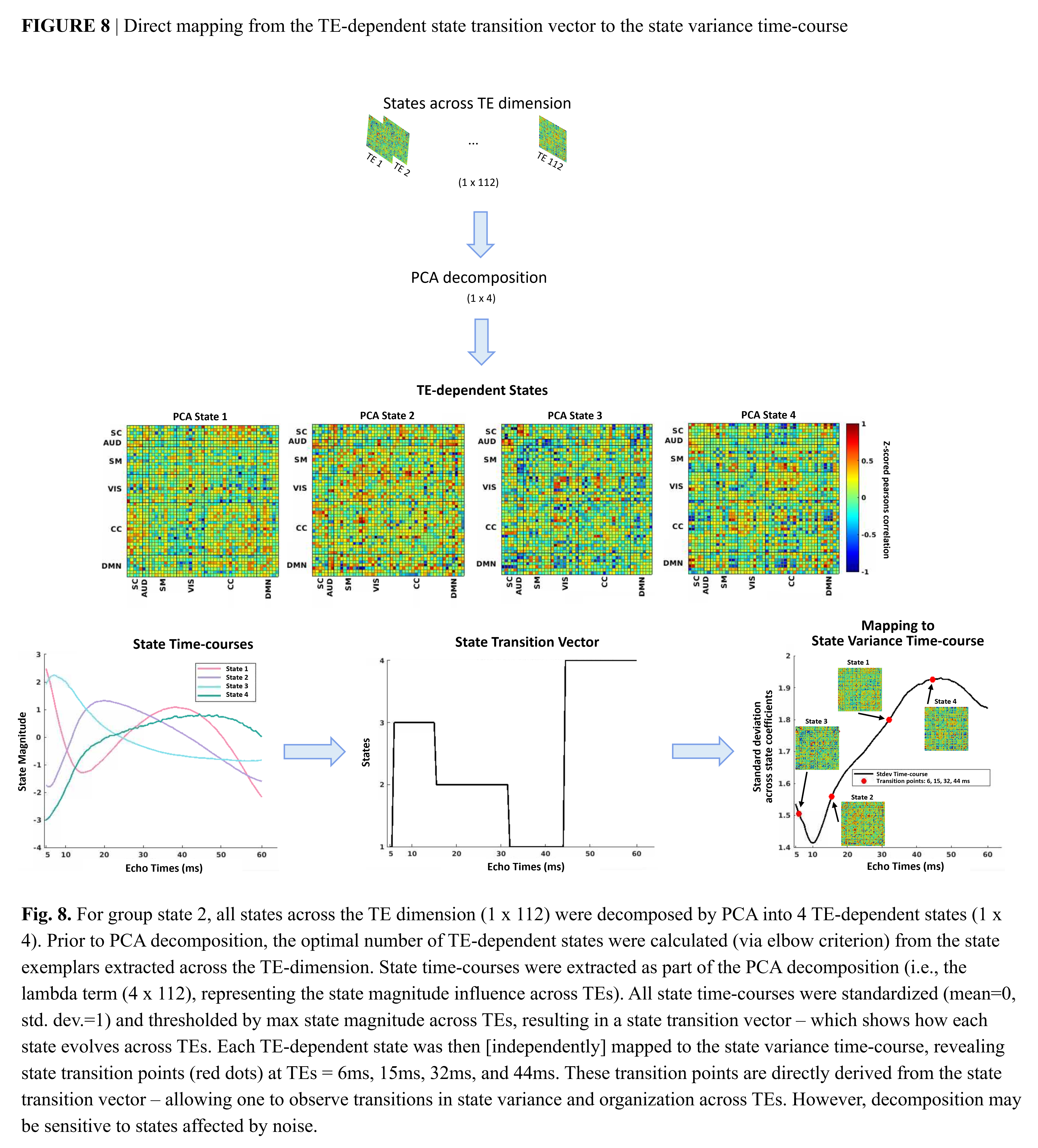
